# Gonadotropes in medaka grow long extensions with varicosity-like swellings, projecting towards each other and blood vessels

**DOI:** 10.1101/777151

**Authors:** Heidi Kristine Grønlien, Romain Fontaine, Kjetil Hodne, Isabelle Tysseng, Eirill Ager-Wick, Finn-Arne Weltzien, Trude Marie Haug

## Abstract

Accumulating evidence in the scientific literature indicates that some pituitary cell types are organized in complex networks. Previous observations have indicated that this may also be the case in medaka (*Oryzias latipes*), where long cellular extensions with varicosity-like swellings are formed by luteinizing hormone (Lh)-producing gonadotropes expressing green fluorescent protein. In this study, immunofluorescence of intact pituitaries reveal that Lh beta polypeptides are mainly located in the varicosity-like swellings and at the extremity of the extensions. Some extensions approach nearby Lh-producing cells, and other extensions are in close contact with blood vessels. To investigate whether these extensions may contribute to network formation, we followed their development using confocal and fluorescent microscopy on primary cultures. During the first two days in culture, the extensions initiated the formation of homotypic cellular networks and clustering. The extensions were classified as either major or minor. Major extensions were several cell diameters long, dependent on microtubules, and displaying varicosity-like swellings at regular intervals. Minor extensions typically protruded from the major, were significantly shorter and thinner, and dependent on actin. The swellings were dependent on both microtubules and actin. Flash photolysis of caged Ca^2+^ showed that the signal was propagated along the major extensions, intensifying in each swelling, indicating a continuous structure. However, the Ca^2+^ signal did not transfer to the next cell in the network, but was transferred between cells merged at their somas. In summary, Lh-producing gonadotropes in medaka display a complex cellular structure of extensions, possibly linked to communication with blood vessels and/or other gonadotrope cells.

## 1. Introduction

The anterior pituitary is a complex endocrine gland that secretes multiple hormones to control lactation, growth, reproduction, and homeostasis. Accumulating evidence indicates that in both mammals and teleost fish, some pituitary cell types are organized into complex three-dimensional homotypic and heterotypic networks (Bonnefont et al., 2005; Budry et al., 2011; Edwards et al., 2017; Golan et al., 2016a; Golan et al., 2016b; Mollard et al., 2012). In teleost fish, the different endocrine cell types are clustered (Golan et al., 2014; Levavi-Sivan et al., 2010; Weltzien et al., 2004), whereas in mammalian pituitaries the different pituitary cell types are distributed in a more mosaic-like pattern (Musumeci et al., 2015).

The anterior pituitary is under control of releasing and inhibiting factors from the brain; in mammals via the portal blood vessel system, in teleosts by axons extending to the pituitary target cells. The neuropeptide gonadotropin-releasing hormone (GnRH) is released in response to certain internal and external cues, and, in turn, stimulates pituitary gonadotropes to express and secrete gonadotropins; follicle-stimulating hormone (FSH) and luteinizing hormone (LH). Downstream, the gonadotropins stimulate gametogenesis and steroidogenesis in the gonads (Weltzien et al., 2004). In mammals and birds, FSH and LH are synthesized and secreted from the same cell (Pope et al., 2006), whereas in most teleost fish, they are produced by distinct cells (Kanda et al., 2011; Weltzien et al., 2014). In teleosts, both gonadotropes are clustered in proximal *pars distalis* (Levavi-Sivan et al., 2010; Weltzien et al., 2004; Zohar et al., 2010). Despite the differences between mammalian and teleost pituitary architecture, teleost pituitary cells are important for comparative studies of neuroendocrine regulation and intracellular signaling pathways (Chang and Pemberton, 2018).

In transgenic medaka (*Oryzias latipes*) that express green fluorescent protein (Gfp) under the control of the endogenous *lhb* promotor (tg(*lhb*:hrGfpII); see (Fontaine et al., 2019; Hildahl et al., 2012), Lh-cells form long extensions in primary culture (Strandabo et al., 2013), and these differ from those previously described in gonadotrope cells from teleosts (Golan et al., 2016b; Hodne et al., 2013) and mammals (Alim et al., 2012; Childs, 1985; Navratil et al., 2007). This observation prompted us to investigate these extensions in more detail.

Evidence is accumulating that a large variety of cells, not only neurons, form extensions that communicate with other cells by direct contact (Kornberg and Roy, 2014). The connection between two cells can be either an open-ended connection through membrane fusion or electrical coupling via gap junctions, or a close-ended connections in which signals or cargo will have to cross two plasma membranes (Eom et al., 2015; Galkina et al., 2013; Kornberg, 2017; Roy et al., 2011; Sherer, 2013). Also in the pituitary, several endocrine cell types including gonadotropes, have been shown to form various extensions (Fontaine et al. 2019 submitted). For teleosts, the extent and the putative roles of such extensions are barely described so far. A plethora of different names has been suggested for structures that may later be shown to be identical or similar. In this article, we have therefore chosen the neutral terms major and minor extensions.

Here we describe and define the nature of the extensions formed from Lh-producing gonadotropes in medaka, both in intact pituitaries and dissociated primary cell cultures. In intact pituitaries we show how the extensions seem to form networks and reach out to blood vessels. In cell cultures, we have investigated dynamic properties not feasible to assess in intact tissue, like extension growth and initiation of contact.

## 2. Materials and metthods

### 2.1 Experimental model

Japanese medaka (*Oryzias latipes*) of the d-rR strain were kept in recirculating systems with water temperature between 25 and 28 °C, salinity (800 μS), and pH (7.6), and light-dark cycle of L14:D10. Fish were fed three times daily on a combination of dry feed (Scientific Fish Food, Special Diets Service, Essex, UK) and newly hatched brine shrimp, *Artemia sp.* (Argent, Redmond, WA, USA). Tg(*lhb*:hrGfpII) medaka, in which expression of Gfp is controlled by the endogenous *lhb* promotor (Hildahl et al., 2012), were used in all experiments. Fish used in the experiments were one-year old and female, except for imaging of intact pituitaries where both sexes where studied. Handling and use of fish were in accordance with guidelines of the Animal Welfare Committee of the University of Oslo. A general permission to keep the animals in the facilities was given by the Norwegian animal research authority (S-2008/108215) and all animals were kept and handled in agreement with their requirements. A specific approval for this study was not needed, as the animals themselves were not experimentally treated (Norwegian legislation for use of animals in research, Chapter II, §6).

### 2.2 Preparation and confocal imaging of pituitary glands

Pituitaries were removed rapidly from the skulls of anesthetized, decapitated medaka and embedded in a drop of 2% low-melting point agarose (Sigma-Aldrich, St. Lois, MO, USA) in the grooves (5 mm in diameter) of cavity microscope slides. The agarose was dissolved in artificial extracellular solution (ECS) containing 134 mM NaCl, 2.9 mM KCl, 2.1 mM CaCl_2_, 1.2 mM MgCl_2_, 0.1% bovine serum albumin (BSA), 10 mM HEPES, and 1.8 mM glucose. The pH was adjusted to 7.75 with NaOH, and the osmolarity adjusted to 280 mOsm using mannitol. Once the agarose had solidified, ECS with a temperature of 26 °C was added on top to keep the agarose moist. A total of 21 pituitary glands (from 14 female and 7 male fish) were examined.

To visualize blood vessels, lipophilic carbocyanine dye (DiI) (Invitrogen, Carlsbad, CA, USA), diluted in 4% paraformaldehyde (PFA) in PBS, was introduced into anesthetized fish by cardiac perfusion, as described in detail previously (Fontaine and Weltzien, 2019). Brain and pituitary were subsequently dissected and fixed overnight at 4 °C in 4% PFA. Tissues were then rinsed with PBS-Tween (PBST), embedded in 3 % agarose, and sectioned with a vibratome (Leica, Wetzlar, Germany). Sections were stained with 4’6-diamidino-2-phenylindole dihydrochloride (DAPI) (1/1000, Sigma-Aldrich) for 20 min and mounted using vectashield (Vector Laboratories, Peterborough, UK).

Whole pituitaries and pituitary sections were imaged using an Olympus FluoView1000 inverted IX81 confocal laser-scanning microscope (Olympus, Tokyo, Japan) with water immersion 4X, 20X, and 40X, or oil 60X (1.10 N.A. Plan APO) objectives as appropriate, or Zeiss LSM710 confocal microscope (Carl Zeiss AG, Oberkochen, Germany) with 63X objective (1.3 N.A. LCI Plan-Neofluar). Optical sections were made every 0.2-0.5 μm along the z-axis and images were obtained through projections of the z-stack. For all pituitaries investigated, the resulting z-stack was studied to ensure that structures that resembled extensions in one section were not the edge of a neighboring cell. Such structures with no cell soma in the section directly above or below, was registered as true extensions.

### 2.3 Tamra dye tracking in pituitary slices

Dye tracing using Tamra (NHS-rhodamine (5/6-carboxy-tetramethyl-rhodamine succinimidyl ester) (ThermoFisher Scientific, Waltham, MA, USA) mixed isomer was conducted using the whole cell patch-clamp technique on brain pituitary slices. Following removal of intact brain with pituitary, 150 μm slices were cut using a vibratome (VT 1000, Leica). Fresh slices were transferred to a recording chamber with ECS. Patch electrodes (borosilicate glass with filament, resistance 5 MΩ) were filled with intracellular solution (mM): 120 CH3SO3K, 20 KCl, 10 HEPES/NaOH, 20 sucrose (pH 7.2), 290 mOsm. Immediately before conducting the experiments, 10 μl Tamra (1mg/ml) was added to 1 ml intracellular solution. Following gigaseal formation, gentle suction was applied to allow complete access of the dye to the whole cell. The cells were visualized using an infrared Dodt Gradient Contrast (DGC) system coupled to an upright Olympus fluorescence microscope with 40X water-immersion objective (0.8 N.A. Slicescope, Scientifica, Uckfield, UK). For excitation of Tamra, a diode light source was used (pE-4000 coolLED, Andover, UK). Tamra was excited at 550 nm, and the emission was collected after passing a 630/75 bandpass filter (Chroma Technology Corp, Vermont, USA). The cells were imaged using a sCMOS camera (optiMOS, QImaging, Surrey, BC, Canada) and both the light source and camera were controlled by μManger software, version 1.4 (Edelstein et al., 2014).

### 2.4 Primary pituitary cell cultures

The cell cultures were made as previously described (Ager-Wick et al., 2018; Strandabo et al., 2013). In brief, the fish were anesthetized, then the pituitary was dissected out and enzymatically digested with trypsin (2 mg/mL, Sigma-Aldrich) for 30 min at 26 °C, followed by incubation with trypsin inhibitor (1 mg/mL, Sigma-Aldrich) and Dnase I Type IV (2 μg/mL, Sigma-Aldrich) for 20 min at 26 °C with gentle shaking. Pituitaries were then mechanically dissociated using a glass pipette, centrifuged at 100 *g* and resuspended in growth medium (L-15, Life Technologies) added 10 mM NaHCO_3_ (in order to stabilize pH at 7.75), 1.8 mM glucose, penicillin/streptomycin (5 000 U per 100 mL medium, Lonza, Verviers, Belgium) and adjusted to 280-290 mOsm with mannitol. Dissociated cells were plated on poly-L-lysine pre-coated dishes fitted with a central glass bottom (MatTek Corporation, Ashland, MA, USA). Each pituitary resulted in a total of 5 000-10 000 cells, and each dish contained about 50 000-100 000 cells, limited to the glass region of the dish. Each culture contained 40-50 pituitaries, divided into four dishes. The cells were incubated at 26 °C and 1 % CO_2_, and used in experiments for up to 5 days after seeding (Ager-Wick et al., 2018).

### 2.5 Fluorescence imaging of primary cell cultures

Culture dishes were imaged using Olympus BX61WI and Zeiss LSM710 confocal microscopes. To study the initial plasticity of Lh-cells and their extensions, time-lapse confocal imaging was conducted immediately after seeding. To determine the cytoskeleton architecture of extensions, cells were exposed to either an inhibitor of actin filaments, cytochalasin B (Sigma-Aldrich), or to a tubulin-polymerization inhibitor, nocodazole (Sigma-Aldrich). Two hours after plating, cytochalasin B or nocodazole [1:5000 dilution from a 10^−2^ M stock in DMSO] was added, giving a final concentration of 10^−7^ M. Control cells were incubated in medium without DMSO as a 1:1000 dilution of DMSO had no effect on extension growth (data not shown). The cells were incubated at RT with cytochalasin B and nocodazole for 2 h followed by live imaging for 4 h. To investigate the effect of Gnrh on extension growth, Gnrh1 (Bachem, Budendorf, Switzerland) was added to the growth medium one day after plating to give a final concentration of 10^−7^ M. The cells were incubated with Gnrh1 for 20 h prior to imaging.

### 2.6 Uncaging of Ca^2+^ in primary cell cultures

Primary pituitary cells were incubated for 1 h at 27 °C with 0.01 % pluronic, 5 μM o-Nitrophenyl EGTA, AM (NP-EGTA-AM, Ca^2+^ caging compound, Thermo Fisher Scientific) and 5 μM Cal-590-AM (Ca^2+^-sensitive dye, AAT Bioques, Sunnyvale, CA, USA) in ECS without BSA, followed by 20 min desertification of NP-EGTA and Cal590 in ECS containing 0.1 % BSA. The cells were visualized as described for Tamra dye tracking. Gfp was excited at 470 nm, and emission was collected after passing a 525/50 bandpass filter (Chroma Systems Solutions). Cal590 was excited at 580 nm and images were collected after passing a 630/75m bandpass filter (ET630/75m emitter, Chroma System Solutions). For Ca^2+^ imaging, an exposure time of between 50 and 80 ms was used and images were sampled at 0.5 Hz. To uncage NP-EGTA, pulses of 50-250 ms were delivered using a 405 nm laser pre-set at 7 mW (Laser Applied Stimulation and Uncaging system, Scientifica). The laser passed through an 80/20 beam splitter and was targeted to distinct regions of the cells using two galvanometer scan mirrors controlled by software (Scientifica) developed in LabVIEW (National instruments, Austin, TX, USA). Images containing laser reflection were removed from the final imaging profile post recording. The relative fluorescence intensity was calculated after background subtraction as changes in fluorescence (ΔF) divided by average intensity of the first 15 frames (F).

### 2.7 Immunofluorescence in primary cell cultures

Two days after plating, the cells were fixed in 4 % PFA for 10 min at room RT, rinsed in PBST several times, and incubated in blocking buffer (10% normal goat serum (Sigma-Aldrich) in PBST) for 1 h. Cells were then incubated with antibody (from rabbit, produced in house) against medaka Lh-beta polypeptides (Lhβ) overnight at 4 °C (Burow et al., 2018), washed in PBST three times, and incubated for 1 h with secondary antibody conjugated to Alexa-fluor 488 (1/1000, Invitrogen) at RT. Finally, the cell culture was washed again in PBST several times before imaging as described above.

### 2.8 Image analysis and processing

Images were processed using the open-source software from ImageJ (versions 1.37v and 2.0.0, National Institute of Health, Bethesda, MD, USA) and Imaris 9.1 (Bitplane, Oxford Instruments Company, Abingdon, UK). Extension length was analyzed with plugin NeuronJ (Meijering et al., 2004). The lengths of major extensions were measured from soma to tip. For intercellular bridges the length was determined as half the bridge length. For branched extensions the length of the secondary extension was measured from the intersection. The smallest diameter was used for analysis.

All photomontages were made in Adobe Photoshop, Illustrator, and Indesign CC2018 (Adobe systems Inc., San Jose, CA, USA).

### 2.9 Quantification and statistical analysis

Numerical data are presented as percentage, mean ± standard deviation (SD) or median and interquartile range (IQR). All data from intact pituitaries are from at least three individuals. All data from primary cell cultures are from at least three independent cultures. In all cases, data distribution was tested for normality. Independent sample t-tests were performed for comparisons of length, diameter and number of extensions, and Mann-Whitney U-tests were performed for comparison of time. The relationship between variables was investigated using Pearson product moment correlation coefficient (r). Preliminary analyses were performed to ensure no violation of the assumptions. *p* values below 0.05 were defined as statistically significant. All analyses were performed using SPSS 23 for Windows (IBM, Armonk, NY, USA). Boxplots were generated by the application BoxPlotR (Spitzer et al., 2014).

## 3. Results

### 3.1 Characteristics of Lh-cell extensions in intact pituitaries

The endogenously produced Gfp in the transgenic tg(*lhb*-hrGfpII) medaka line provides a robust fluorescent signal from all parts of the cell, also during prolonged confocal imaging, and thus enabled extensive and detailed mapping of anatomical structures within the tissue. Confocal images from intact pituitaries revealed that Lh-cells form numerous long extensions (Fig. 1A). Interestingly, the extensions displayed multiple distinct areas of enlargement, hereafter referred to as swellings (arrows). Similar structures were observed in pituitaries from both male (n=7) and female (n=14) fish. Furthermore, filling cells in pituitary slices with Tamra repeatedly displayed long cellular extensions with varicosity-like swellings (n=5, Fig. 1B). Tamra was never transferred to other cells in contact with the filled one.

**Fig. 1.**
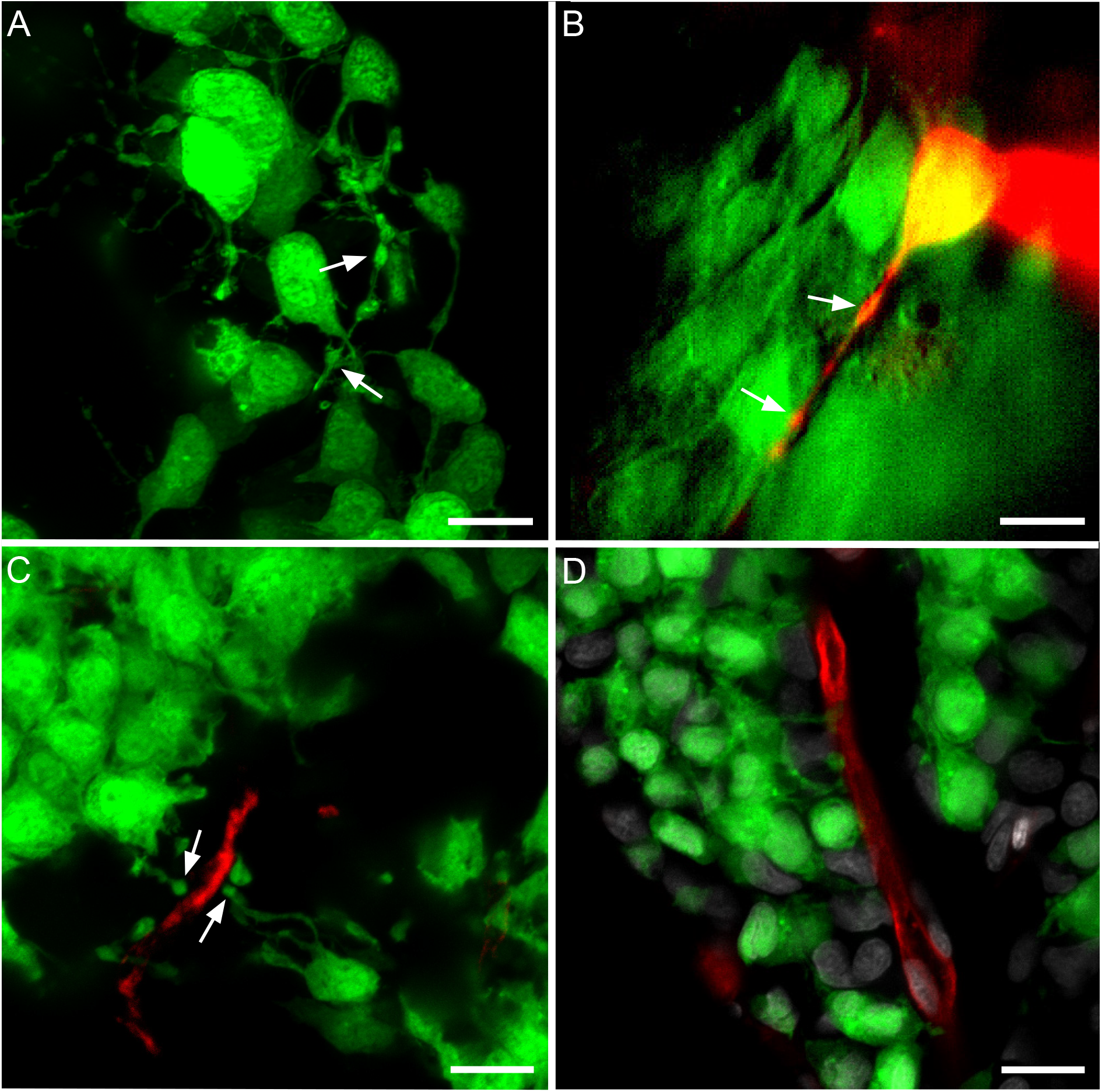
Characteristics of Lh-cell extensions in whole pituitary. (A) Projection of 10 μm confocal z-stack of Lh-cells in whole pituitary. Arrows show swellings. Fluorescence image of a single Lh-cell in pituitary slice with Tamra dye (red) injected through a patch pipette. Arrows show swellings. (C) Projections of 10 μm confocal z-stack of Lh-cells in a fixed pituitary section with DiI cardiac perfusion in advance of dissection. Arrows show swellings nearby a blood vessel. (D) Projections of 10 μm confocal z-stack of Lh-cells in a fixed pituitary section with DiI cardiac perfusion in advance of dissection, with nuclear DAPI staining. Note that extensions from Lh-cells project towards the blood vessel. Scale bars represent 10 μm.

Pituitary sections after cardiac perfusion with DiI, which labels the endothelium, showed that some Lh-cells are organized around blood vessels, with their extensions projecting towards the vessel (Figs. 1C and D). The terminal part of the extension appeared wider where it reached the blood vessel (Fig. 1C).

### 3.2 Characteristics of the Lh-cell extensions in primary cell cultures

#### 3.2.1 Extensions can be divided in two categories: major and minor

During dissociation, the cells lost all their extensions. However, after approximately 1 h (median 66 minutes, IQR 128 min; n=45 cells), the Lh-cells started forming new extensions. After 2 days, extensions with several swellings were observed from 99 % of the Lh-cells (Fig. 2). The overall number and length of the extensions plateaued after 2 days, therefore structural analyses were conducted at this time point. Extensions were also observed from about 20 ± 10 % (n=3 cultures, in total 517 cells) of the non-*lhb*-expressing cells, but these were not investigated further.

**Fig. 2.**
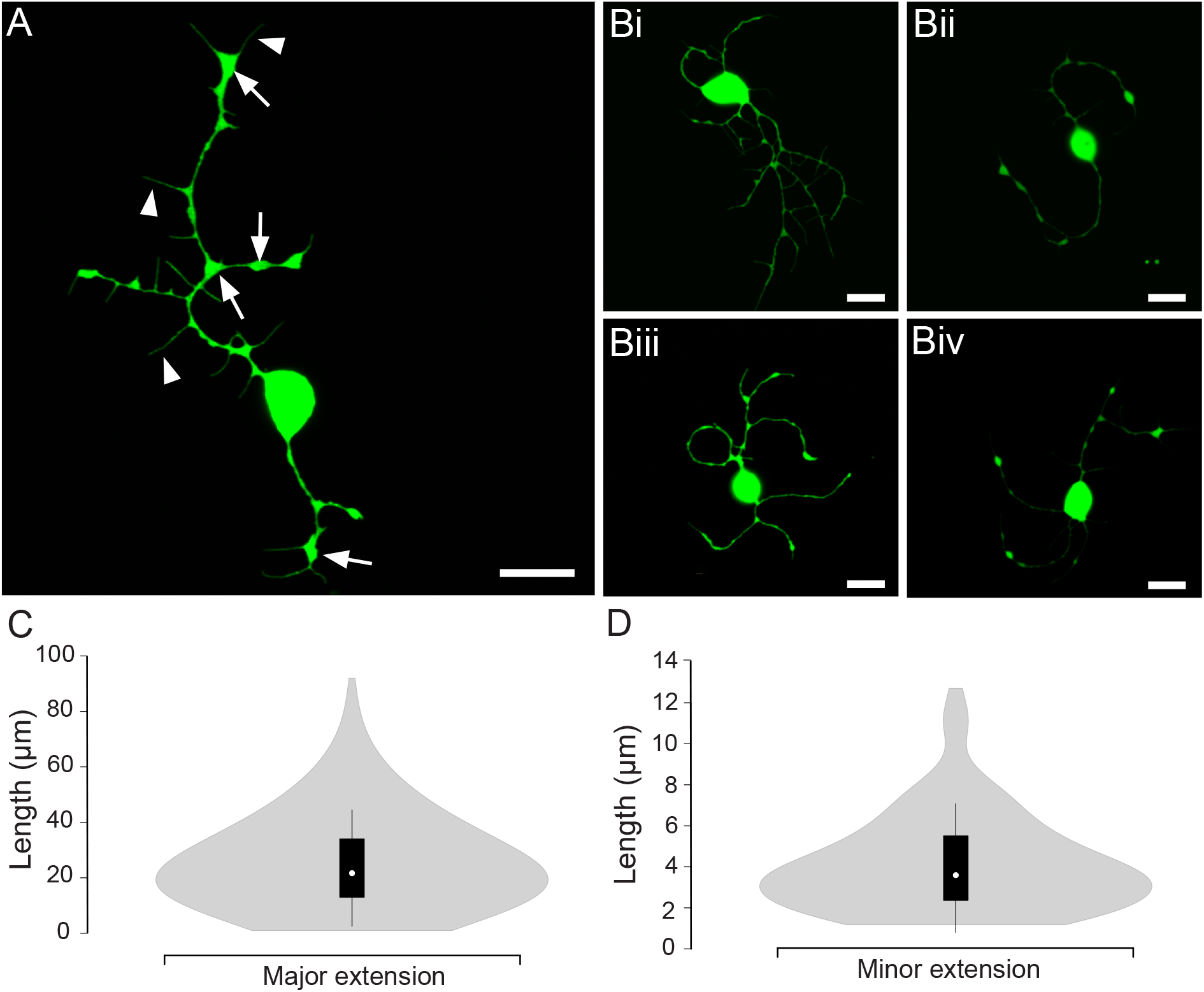
Characteristics of Lh-cells with major and minor extensions in primary cell culture. (A) Projection of a confocal plane of an Lh-cell from two days old dissociated primary pituitary cell culture. Arrows show swellings and arrowheads show minor extensions. (Bi-iv) Set of planar confocal projections of Lh-cells from primary cell cultures. Images present the heterogeneity of the Lh-cells. Scale bars represent 10 μm. Violin plots of major extension length from 741 extensions (C) and minor extension length from 114 extensions (D). White circles show the medians; box limits indicate the 25th and 75th percentiles as determined by R software; whiskers extend 1.5 times the interquartile range from the 25th and 75th percentiles; polygons represent density estimates of data and extend to extreme values.

The major extensions were heterogenic in appearance and size (Table 1, Figs. 2A and B). Figure 2C presents the distribution of length of 741 extensions from 311 Lh-cells. 90% of the extensions of Lh-cells displayed swellings; these were regularly spaced, with mean distance 10.9 ± 5.8 μm (n=60 cells/103 extensions). The cross-sectional area of a swelling constituted 6.8 ± 3.8 % of the area of the soma (n=60 cells/103 extensions). For approximately 75% of the Lh-cells (n=232 cells), one or more of the major extensions branched.

**Table 1.** Characteristics of major and minor extensions of Lh-cells in primary cell cultures at 48 h.

|  | **Major extension** | **Minor extension** |
| --- | --- | --- |
| *Diameter (mean ± SD)* | 0.87 ± 0.28 μm (n=158) | 0.24 ± 0.05 (n=114) |
| *Length (mean ± SD)* | 25.1 ± 15.6 μm (n=741) | 3.4 ± 2.3 (n=114) |
| *Appearance* | Irregular, bending | Uniform, straight |
| *Cytoskeleton* | Microtubules and<br>actin filaments | Actin filaments |
| *Number of extensions*<br>*(mean ± SD)* | 2.5 ± 1.3 (n=311 cells) | 6.9 ± 3.4 (n=16 major extensions) |
| *Swellings* | Yes | No |
| *Branching* | Yes | No |

Minor extensions were growing from all regions of the major extensions, including the tip and swellings (Fig. 2A). For each major extension, 6.9 ± 3.4 (n=16 major extension) minor extensions were observed. These minor extensions were thinner, shorter and uniform in thickness without swellings (Table 1). The number of minor extensions correlated with the length of their major extension (Pearson’s correlation coefficient (r) of 0.6, n=16, *p*=0.015). Figure 2D shows the distribution of length of 114 minor extensions from 9 Lh-cells; none of the minor extensions exceeded 13 μm in length.

#### 3.2.2 Ca^2+^ signals propagate from Lh-cell soma to the terminal

To examine if the extensions are continuous structures where Ca^2+^ signals can propagate through the extensions, Ca^2+^ was uncaged in the soma of an Lh-cell. Monitoring the cytosolic Ca^2+^ concentration showed clearly that the Ca^2+^ signal propagated along the extension from the soma to the swellings, reaching the terminal of the cell (Fig. 3 and supplementary video S1).

**Fig. 3.**
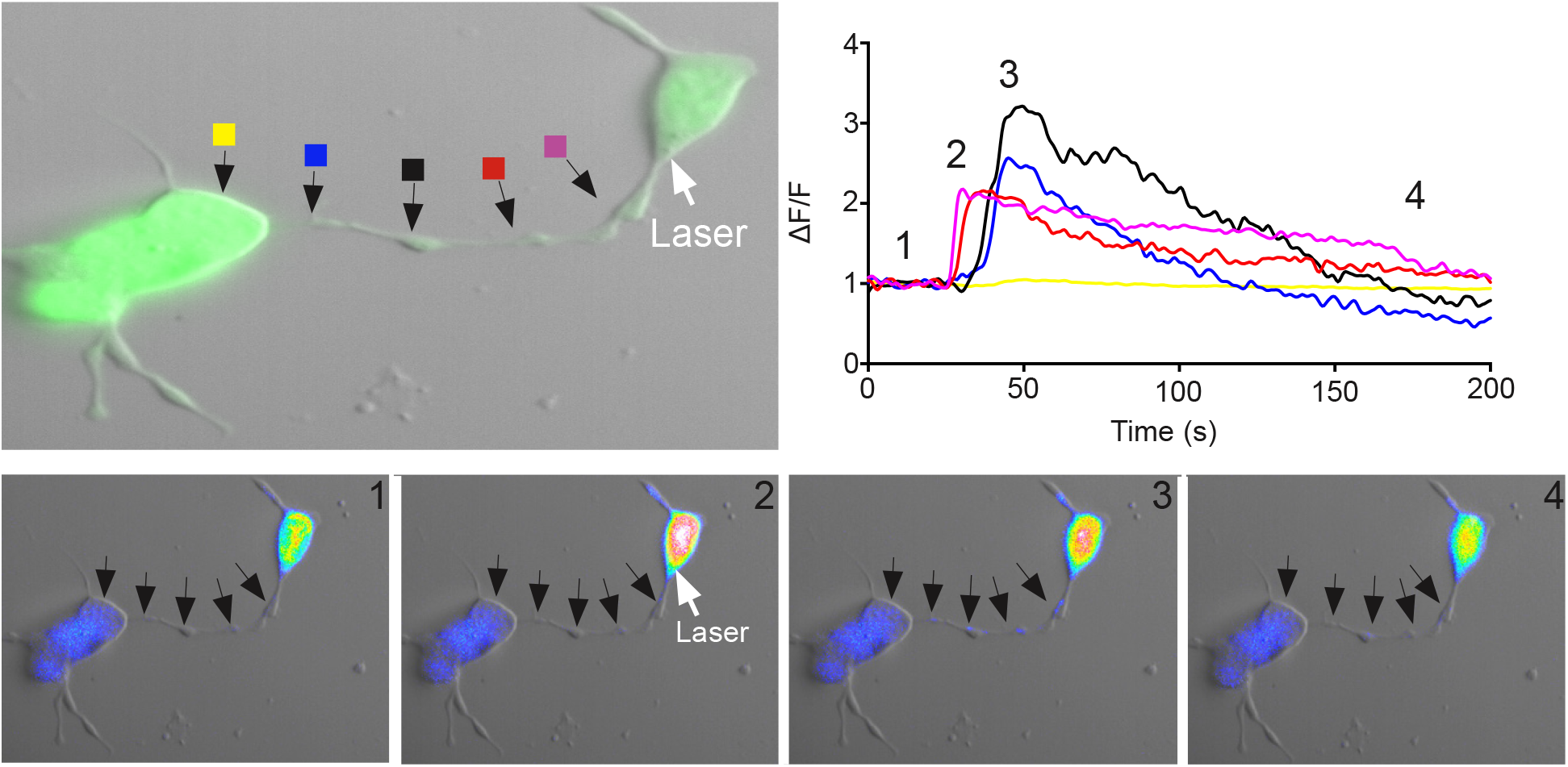
Ca^2+^ signal propagation in an extension from an Lh-cell in primary cell culture. Flash photolysis of caged Ca^2+^ in cultured Lh-cells recorded by Ca^2+^ imaging. Upper left panel shows two Lh-cells connected by an extension. The area of Ca^2+^ uncaging is marked with an arrow; colored boxes indicate the sites of intracellular Ca^2+^ measurement shown in the upper right panel. Changes in fluorescence (ΔF) divided by the average intensity of the first 15 frames (F) as a function of time. The pictures in the lower panels show imaging of intracellular Ca^2+^ concentration at the four time points indicated in the upper right graph. Note that the Ca^2+^ signal propagates along the extension, but does not transfer to the connected cell. See also video S1.

#### 3.2.3 Lh-cell extensions contain Lhβ protein

Although the previous figures clearly show that the extensions and swellings are filled with GFP, this does not imply that the Lh hormone is located here, as the GFP protein is not linked to the Lhβ protein. To visualize the location and distribution of the hormone itself, immunofluorescent labeling of the Lhβ protein was performed. High fluorescence was detected in the major extensions, with the swellings and the extremities of the extensions more strongly labelled than the cell soma (Fig. 4), suggesting a high quantity of Lhβ protein in these cellular regions. Although the immunofluorescent labelling is presented in green, it should thus not be confused with the endogenously produced GFP.

**Fig. 4.**
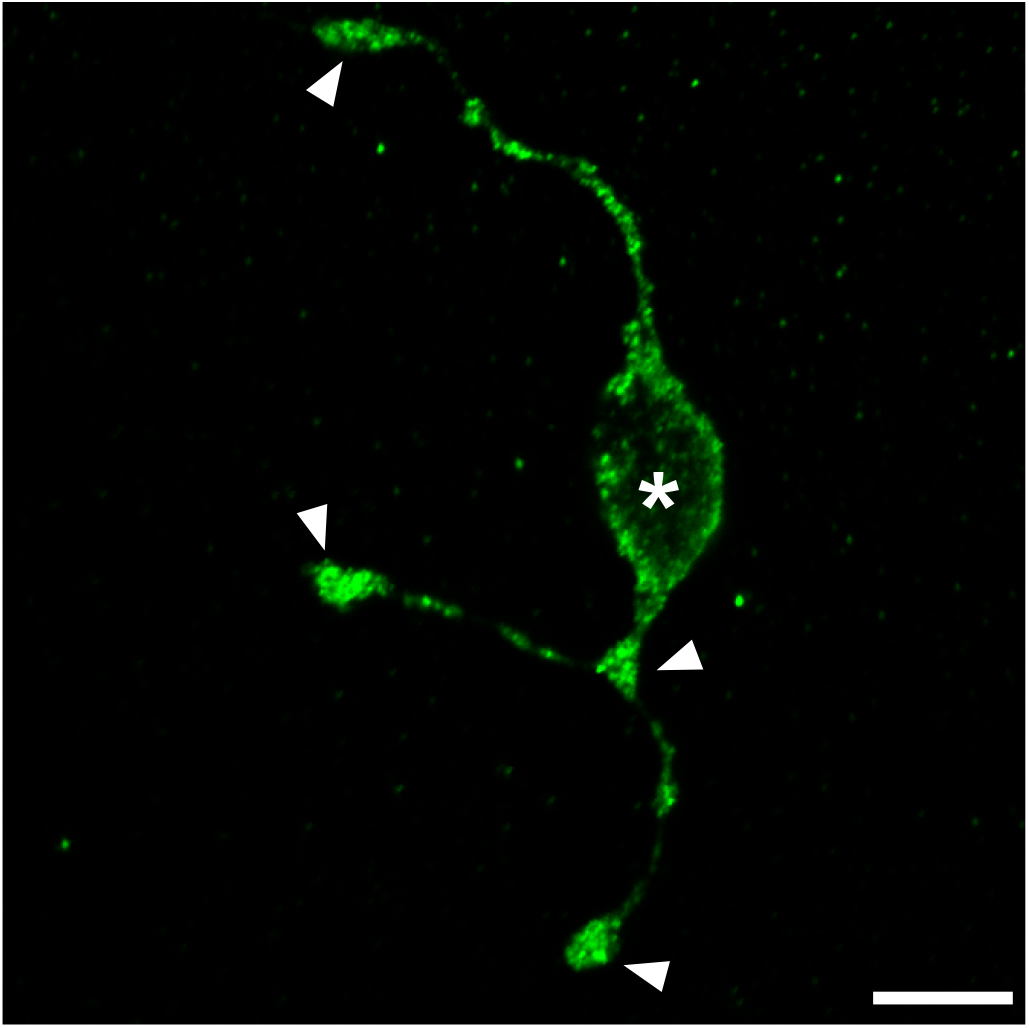
Lhβ proteins in an Lh-cell in primary cell culture. Planar confocal projection of an Lhβ immunoreactive cell. Arrows show the swellings and the extremities of the extensions with high content of Lhβ protein, and asterisk shows the cell body. Scale bar represents 10 μm.

#### 3.2.4 The cytoskeleton architecture of Lh-cell extensions

Following incubation with the tubulin polymerization inhibitor, nocodazole, Lh-cells (n=21) produced only one type of extension (Fig. 5B). These were uniform in thickness, displayed no swellings, fewer than 10 % were branched, and they were thinner than the previously described major extensions, with a mean diameter of 0.29 ± 0.10 μm (n=130) (Fig. 5D). Thus, nocodazole-treated cells did not form typical major extensions. In contrast, Lh-cells (n=27) that had been incubated with cytochalasin B, an inhibitor of actin filaments, produced extensions with a mean diameter of 0.84 ± 0.10 μm (n=84) (Fig. 5D), which is similar to the previously described major extensions, but without the distinctive swellings (Fig. 5C). Of the cytochalasin B-treated cells, approximately 40 % displayed branching extensions, minor extensions were not observed, and only 5 out of 84 extensions were straight. In addition, the somas generally displayed an irregular shape (Fig. 5C).

**Fig. 5.**
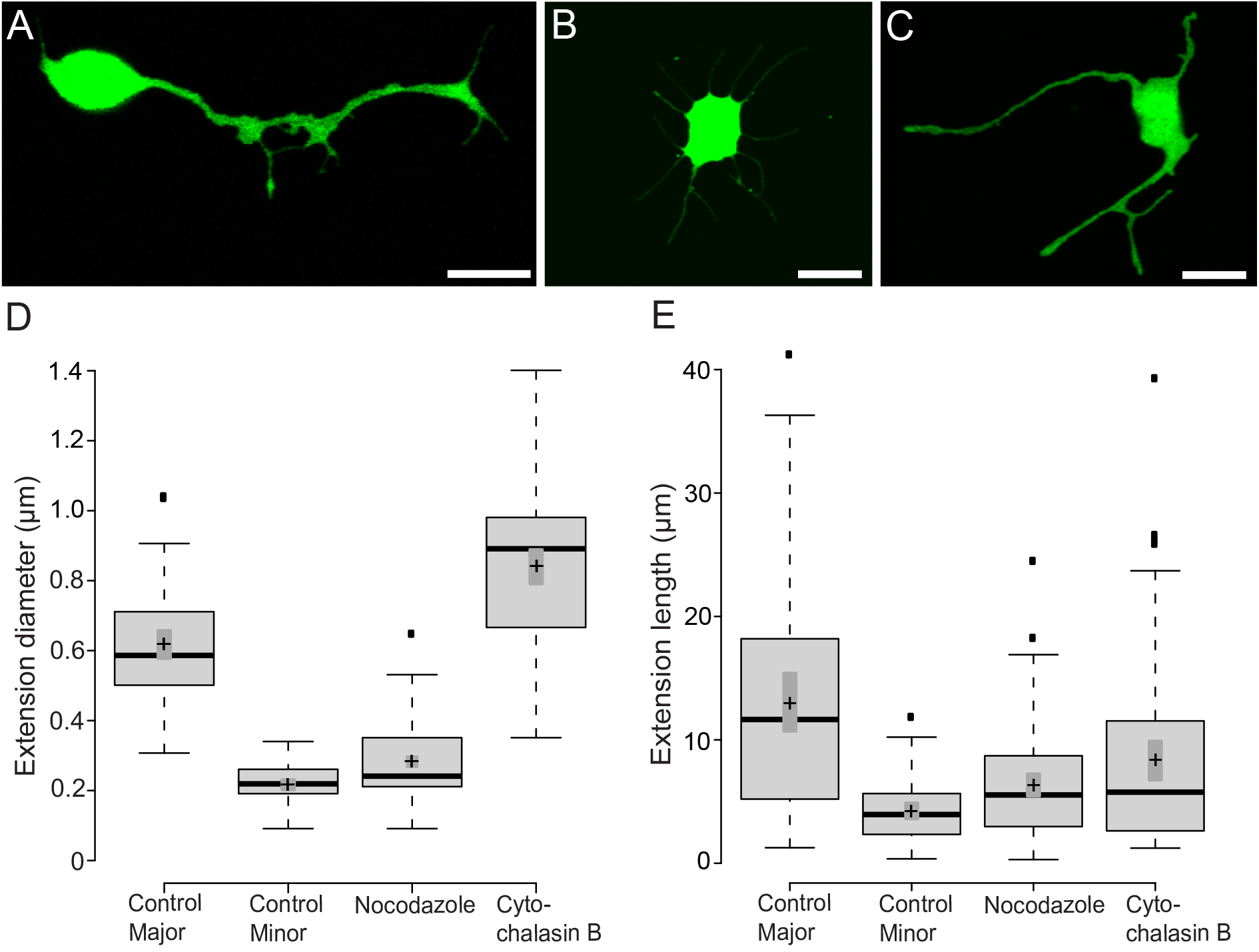
Characteristics of Lh-cell extensions in culture treated with cytoskeleton inhibitors. Planar confocal projections of (A) a non-treated Lh-cell (control), (B) an Lh-cell treated with 10^−7^ M nocodazole, and of (C) an Lh-cell treated with 10^−7^ M cytochalasin B, 6 h after seeding. Scale bars represent 10 μm. (D-E) Boxplot of extension diameter and length 6 h after seeding. Center lines show the medians; box limits indicate the 25th and 75th percentiles as determined by R software; whiskers extend 1.5 times the interquartile range from the 25th and 75th percentiles, outliers are represented by dots; crosses represent sample means; bars indicate 95% confidence intervals of the means. n = 66, 69, 100, 84 extensions. Nocodazole and cytochalasin B are inhibitors of microtubules and actin filaments, respectively.

#### 3.2.5 Morphological effects in Lh-cells following Gnrh1 exposure

To assess whether Gnrh may affect the appearance of extensions, we exposed of one day old primary cultures to 10-7 M Gnrh1, the most commonly used concentration for hypothalamic releasing hormone studies. These cells were thus regularly exposed to Gnrh1 in vivo during their one-year old lifetime, then restricted from the hormone for one day before being reintroduced to Gnrh1 for 20 h. The exposure resulted in 85 % of the exposed cells (n=120) developing an irregularly shaped soma, with a significant higher number of minor extensions (27 ± 13 minor extensions per cell, n=17 cells) compared to unexposed cells (10 ± 6 minor extensions per cell, n=16 cells, p<0.001) (Figure 6). In addition, a significant increase in the total number of major extensions from each cell was observed (p<0.001); in the unexposed controls, the mean number of major extensions per Lh-cell was 3.1 ± 1.6 (n=83 cells), whereas 5.8 ± 3.1 major extensions per Lh-cell were observed in Gnrh1-exposed cultures (n=80 cells).

**Fig. 6.**
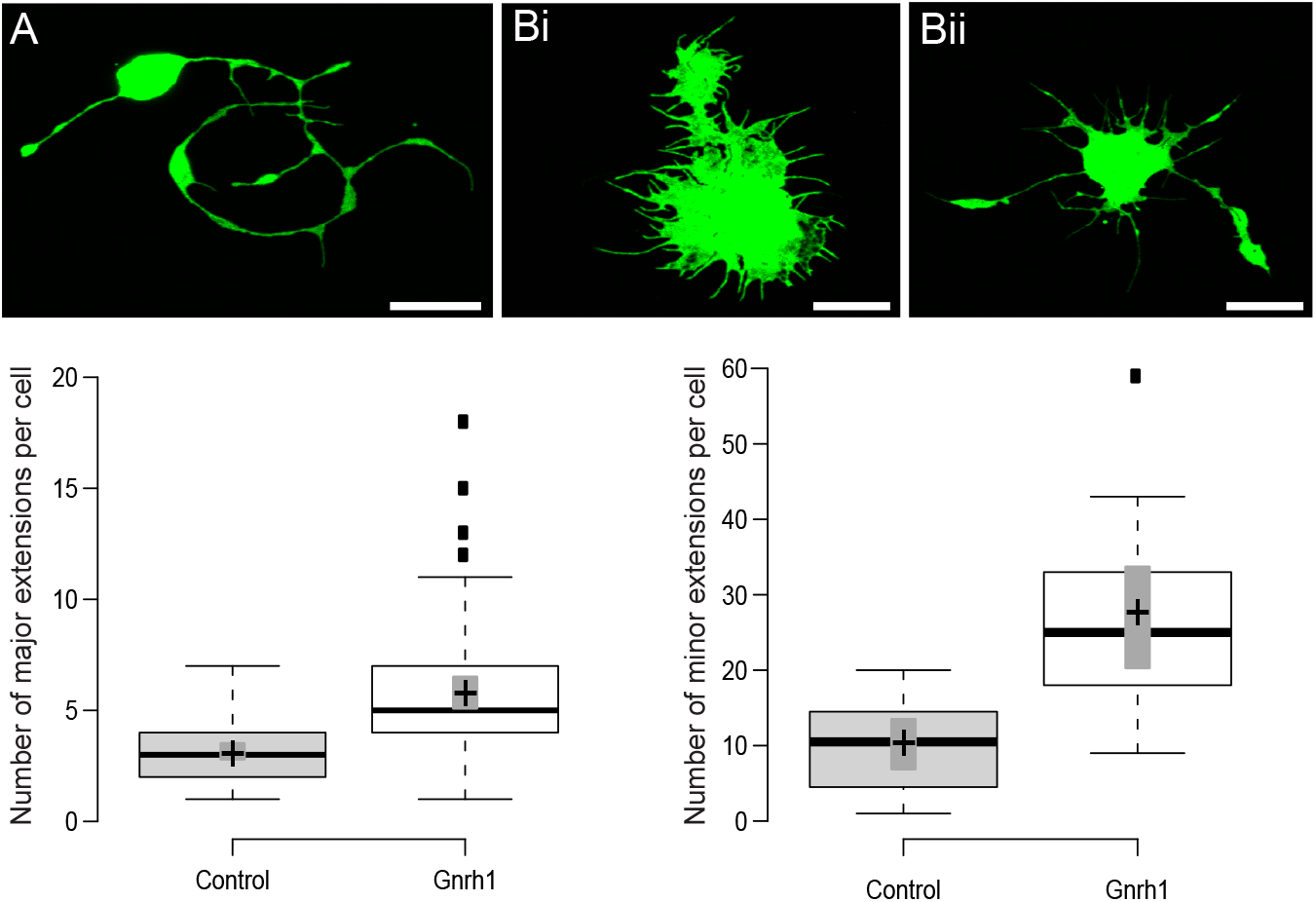
Characteristics of Lh-cell extensions in primary cell culture incubated with Gnrh1. (A) Planar confocal projections of a Lh-cells incubated with 10^−7^ M Gnrh1 for 20 hr. Note the irregularly shaped soma with a number of minor extensions of the Gnrh1 treated Lh-cells. Scale bars represent 10 μm. (B-C) Boxplot of number of major and minor extensions per cell after incubation with 10^−7^ M Gnrh1. Center lines show the medians; box limits indicate the 25th and 75th percentiles as determined by R software; whiskers extend 1.5 times the interquartile range from the 25th and 75th percentiles, outliers are represented by dots; crosses represent sample means; bars indicate 95% confidence intervals of the means. n = 83, 80 cells and n = 16, 17 cells for determining of number of major and minor extensions respectively.

### 3.3 Contact between extensions from different Lh-cells in primary cell cultures

In the intact pituitary, most of the Lh-cells cluster in groups (Fig. 1A), and a similar clustering of Lh-cells was observed in cell cultures (Fig. 7A). To assess whether the extensions play a role in the clustering process, we followed the cells in time-lapse recordings directly after seeding (Fig 7B and supplementary video S2). Of 16 interactions between two Lh-cells, 7 contacts were made by an extension from one Lh-cell connecting with an extension from the other Lh-cell (Fig. 7B). The distance between the somas of the Lh-cells at the point of connection was 27.2 ± 9.8 μm (n=7), and the median time from when the extensions were 10 μm from each other until contact was established was 6.0 min (IQR = 3.0 min, n=7). Contact could also be made directly between an extension and the soma of another Lh-cell (Fig. 7B). Of the 16 interactions analyzed, 9 such contacts were made. Here, the distance between the somas of the two Lh-cells was 13.7 ± 4.7 μm (n=9). Thus, the distance was significantly shorter than when two extensions from different Lh-cells connected (*p*<0.001). The median time from when the extension was 10 μm from the other cell’s soma to connection was made was 12.0 min (IQR=12.0, n=9), which is significantly longer than when two extensions the same distance apart connect (*p*=0.04).

**Fig. 7.**
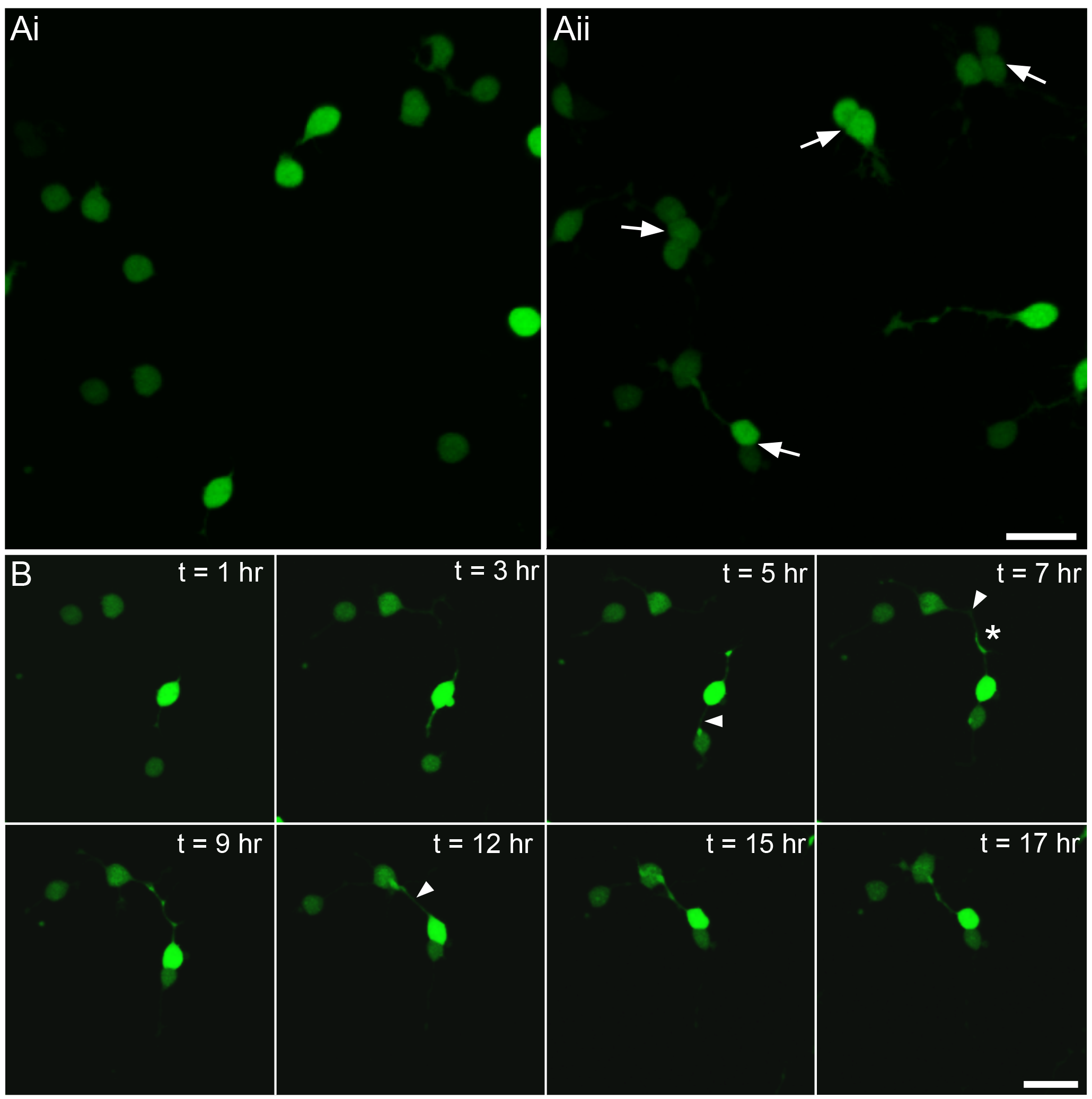
Development of extensions in primary cell culture. (Ai) Planar confocal projections of Lh-cells immediately after seeding and (Aii) 17 h after seeding. Arrows show Lh-cells clustered in groups. (B) Time-lapse projections of four Lh-cells. Time zero is immediately after seeding. Arrowheads show extensions making intercellular contact followed by formation of an intercellular bridge. Asterisk marks a non-Lh-cell. See also video S2. Scale bars represent 10 μm.

Both ways of contact between two Lh-cells, either extension-extension or extension-soma, resulted in formation of a continuous structure without swellings that connected the two somas, hereafter referred to as an intercellular bridge. The bridges were dynamic in nature, following the cells’ movements without breaking (Fig. 7B). The intercellular bridges typically straightened after contact, with median time between contact and straightening of the bridge being 12.0 min (IQR=25.5 min, n=16). Bridge formation was followed by a period in which the Lh-cells moved closer together (Fig. 7B) with a mean shortening distance of 10.4 ± 3.1 μm (n=16). The shortening was no different regardless of whether the bridge had been made by two extensions connecting or by one extension connecting to the soma of another Lh-cell. The diameter of each bridge was 1.05 ± 0.27 μm (n=16), which is significantly thicker than the diameter of a major extension (*p*<0.001). During the first 10 h, most cell-cell connections that formed developed into bridges, followed by clustering soma to soma, and in our 2-day cultures, 40 % of the Lh-cells were clustered (n=623 Lh-cells). Although the Lh-cells tended to form homotypic networks in primary cell cultures, many extensions were not in contact with other cells (Fig. 8).

**Fig. 8.**
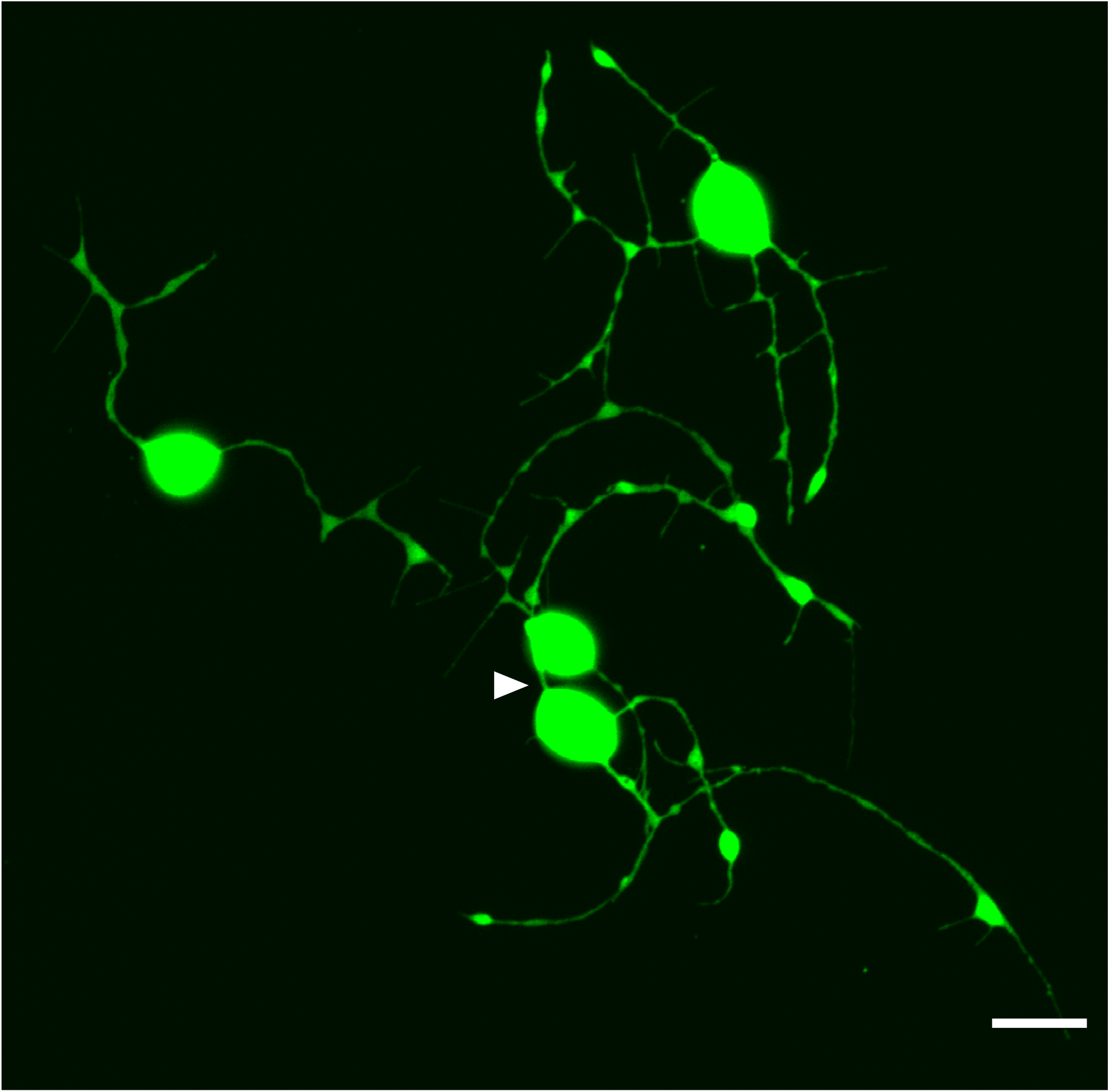
Network of Lh-cells in primary cell culture. Projection of a confocal plane of Lh-cells forming a network in three days old primary culture. Arrowhead shows an intercellular bridge between two Lh-cells. Scale bar represents 10 μm.

As shown earlier (Fig. 3) by monitoring the cytosolic Ca^2+^ concentration after uncaging Ca^2+^ in the soma of one cell, the Ca^2+^ signal propagated to the border between the extension and the cell to which it was connected. However, we did not observe any transfer of Ca^2+^ signal between the cells (n=20), neither via regular extensions nor via the straight intercellular bridges (n=13) (Fig. 9). Occasionally (n=3), we observed an elevation in intracellular Ca^2+^ concentration in extensions that we were unable to assign to neither the stimulated cell nor its neighbor (Fig. 9).

**Fig. 9.**
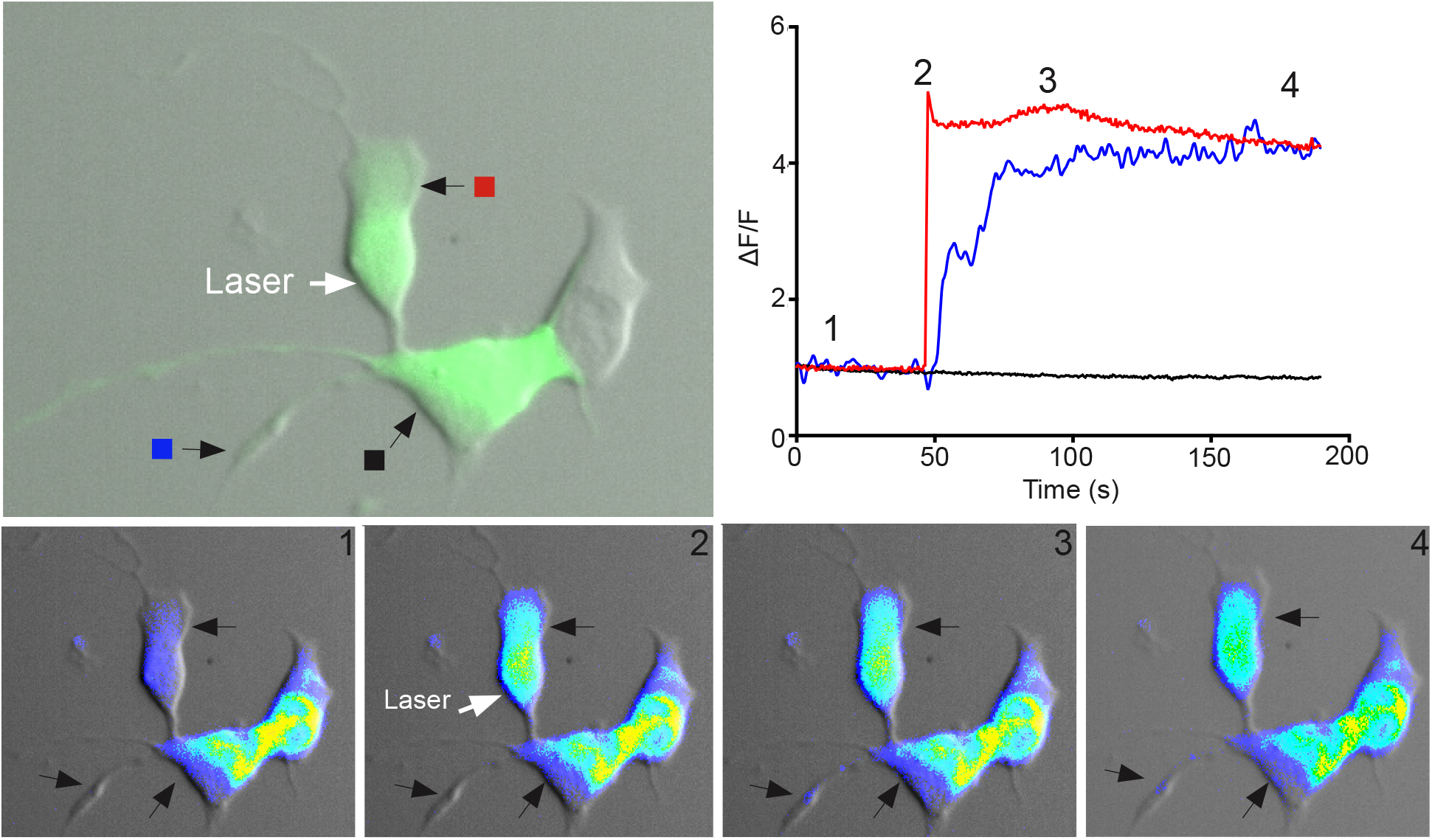
Ca^2+^ signaling between Lh-cells connected soma to soma in primary cell cultures Flash photolysis of caged Ca^2+^in cultured Lh-cells recorded by Ca^2+^ imaging. Upper left panel shows two Lh-cells connected by an intercellular bridge. The upper cell is merged to another Lh-cell, soma to soma. The area of Ca^2+^ uncaging is marked with an arrow; colored boxes indicate the sites of intracellular Ca^2+^ measurement shown in the upper right panel. Changes in fluorescence (ΔF) divided by the average intensity of the first 15 frames (F) as a function of time. The pictures in the lower panels show imaging of intracellular Ca^2+^ concentration at the four time points indicated in the upper right graph. Note that the Ca^2+^ signal transfers across the soma-soma border, but not across the intercellular bridge. See also video S3.

In contrast, when two or more cells were tightly merged soma to soma (Fig. 9 and supplementary video S3), in 71 % of all experiments (n=21), uncaging of Ca^2+^ in the soma of one cell induced a significant elevation of cytosolic Ca^2+^ in one or more of its merged neighbor cells. The elevation in the neighbor cell occurred within 500 ms after uncaging Ca^2+^, and the peak response normally occurred 1-2 s after uncaging.

## 4. Discussion

The established view regarding the morphology of endocrine cells from the anterior pituitary is that they have a uniform, rounded shape with few or no extensions of any type. This view has been repeatedly challenged in recent years, as more refined imaging methods have become available (Golan et al., 2016b; Le Tissier et al., 2016; Le Tissier et al., 2015; Mollard et al., 2012). Our transgenic medaka line, in which the Lh-producing cells are filled with Gfp, display a surprisingly complex pattern of extensions, apparently forming networks with each other. Although these extensions could be suspected to be an artifact associated with the primary culture protocol, our results show that similar structures are also clearly visible in intact pituitary. This spurs several intriguing lines for functional studies to explore the putative roles of pituitary endocrine cell extensions. Some putative roles are suggested in the following discussion, but need to be studied in greater detail.

Cellular extensions in non-neuronal cells have been subject of considerable attention lately (Antanaviciute et al., 2014; Eom et al., 2015; Kornberg and Roy, 2014; Navratil et al., 2007; Rustom et al., 2004). However, as early as 1985, it was noted that mammalian gonadotropes could be stimulated to form extensions by GnRH (Childs, 1985). These findings have been confirmed (Alim et al., 2012; Navratil et al., 2007), but the extensions appear to be structurally quite different from the extensions on medaka Lh-cells. To the best of our knowledge, structures similar to the extensions found on the medaka Lh-cells have not been reported elsewhere. The extensions appear remarkably similar to neurites, regarding diameter, length, dependency on microtubules and the fact that the extension is a continuous structure (Tennyson, 1970; van Beuningen and Hoogenraad, 2016). In addition, they both divide at distinct branching points, and display regularly spaced swellings (Malkinson and Spira, 2010; Tennyson, 1970). The structural similarity of the swellings to axonal varicosities, combined with the fact that they contain Lh, suggests that there may be exocytosis of Lh or other paracrine signaling molecules from these swellings; this would be an interesting topic for future investigation.

The minor extensions of the Lh-cells are dependent on actin and are thus similar to the filopodia that originate from the growth cone of neurites (Gomez and Letourneau, 2014; Smith, 1988). The lack of distinct swellings on the nocodazole- and the cytochalasin B-treated cells suggests that both microtubules and actin filaments are involved in the structure of the swellings, as has been reported for axonal varicosities (Dent and Gertler, 2003). Intermediate filaments have been indicated to play a role in cytoskeletal rearrangements (Leduc and Etienne-Manneville, 2015), and a possible interplay between microfilaments, microtubules, and intermediate filaments in Lh-cell extensions would be worthy of further investigation.

The most striking differences between the processes observed on mammalian gonadotropes and medaka Lh-cells are that in the latter, the extensions are longer, branched, have varicosity-like swellings, and the cytoskeleton is not only actin-based. It is important to note, however, that because cellular extensions are extremely fragile, some of the observed structural differences may reflect artifacts associated with differences in experimental protocols.

There is evidence from mammalian studies that Gnrh induces plasticity of gonadotropes and their extensions by remodeling of the actin cytoskeleton (Edwards et al., 2017; Navratil et al., 2007).We did not aim to extensively characterize such an effect on the Lh cells, but rather confirm that also in medaka, the gonadotrope extensions may be controlled by Gnrh. The 20 h exposure to Gnrh did seem to affect extension growth. In particular, Gnrh seemed to alter the basis of the extensions, making it more similar to a lamellipodium. Lamellipodia are the dynamic domain for cell motility and require organization of actin filaments into lamella networks and bundle arrays (Abercrombie, 1980; Small et al., 2002). One may speculate that a Gnrh-induced increase in connections between Lh-cells during sexual maturation may enhance the efficiency of the coordinated Lh release necessary for reproduction. More elaborate studies are clearly needed to clarify the role of Gnrh in extension growth and Lh cell network formation.

The contact made by extensions during the initial hours after seeding often resulted in an intercellular bridge forming between two Lh-cells. The bridge was apparently strong enough to participate in drawing the two connected cells closer to each other. The ability of filopodia to pull and push have attracted considerable attention (Bornschlögl, 2013), and future research may reveal the cytoskeleton dynamics behind the bridge shortening. In contrast to mammalian gonadotropes that are scattered throughout large parts of the anterior pituitary, teleost gonadotropes are clustered (Levavi-Sivan et al., 2010; Musumeci et al., 2015; Weltzien et al., 2014). This means that there must be a targeted clustering of gonadotropes during development, even separating between Lh-expressing and Fsh-expressing cells, which might be controlled by Gnrh. Godoy and colleagues (Godoy et al., 2011) have suggested that the morphological plasticity of gonadotropes has a developmental role in the organization of the anterior pituitary, as Gnrh induces cell migration in addition to formation of extensions.

Corticotrope cells in the mammalian pituitary use their extensions (cytonemes) to form homotypic and heterotypic networks in addition to promoting long-distance secretion of ACTH (Budry et al., 2011). While in the present study, Lh-cells rarely connected to non-Lh-cells, mammalian gonadotropes form a network that closely follows the pre-established corticotrope network, and cells of these lineages maintain direct contact throughout adulthood (Budry et al., 2011). It is noteworthy that many of the contacts between gonadotropes and corticotropes in mammals involve extensions originating from the corticotropes (Budry et al., 2011; Fontaine et al., submitted; Mollard et al., 2012). A similar heterotypic network is not expected in teleosts, where the gonadotropes and corticotropes are clustered in separate areas. Thus, we would not expect a similar degree of contact between Lh cells and non-Lh cells (i.e. corticotropes) in medaka and other teleosts either.

Ca^2+^ ions play an important role in cell signaling, i.e., for regulating hormone release (Stojilkovic, 2012). The transfer of Ca^2+^ signal from soma to the terminal of an extension, together with the significant concentration of hormone present in the swellings, opens up for the possibility that the extensions may have a role in secretion. In both mammals and teleosts there are indications that gonadotrope extensions may reach out to nearby blood vessels, resulting in more efficient release of hormone into the circulation (Alim et al., 2012; Childs, 1985; Childs et al., 1983; Golan et al., 2016b; Golan et al., 2015; Navratil et al., 2007). Our results indicate that this may also be the case in medaka. We show that the extensions of Lh-cells contain a high quantity of Lhβ protein and project into the perivascular space, offering the possibility of close contact between Lh-cells and the microcirculation.

We observed an extensive network of Lh-cells in both intact pituitary tissue and primary cell cultures, where the extensions seemed to be in physical contact. Ca^2+^ signals were not directly transferred from one cell to another through extensions, and further studies are necessary in order to determine whether the Lh-cells communicate via extensions in other ways, or whether the contacts are merely physical, closed-ended. Previous research has shown that gonadotropes may be functionally linked in larger networks by gap junctions, resulting in efficient spread of regulating signals (Golan et al., 2016b; Gongrich et al., 2016; Levavi-Sivan et al., 2005). The effective transfer of Ca^2+^ signals between merged cells shown in our study may be explained by the presence of gap junctions in the soma-soma connections, but not in the extensions. This also calls for more functional studies in the future.

## Conclusion

Lh-expressing gonadotropes from medaka display neurite-like extensions with varicosity-like swellings containing Lh, both in culture and intact pituitary. Some extensions contact neighboring Lh-cells, apparently inducing clustering or forming Lh-cell networks. Other extensions project towards nearby capillaries.

## Abbreviations

ACTH: Adrenocorticotropic hormone (mammals)
DiI: lipophilic carbocyanine dye
FSH/Fsh: Follocle-stimulating hormone protein (mammals/fish)
GFP: green fluorescent protein
GnRH/Gnrh: Gonadotropin releasing hormone protein (mammals/fish)
IQR: interquartile range
LH/Lh: Luteinizing hormone protein (mammals/fish)
Lhβ/Lhβ: Luteinizing hormone β subunit protein (mammals/fish)
Lhb/lhb: Luteinizing hormone β subunit gene (mammals/fish)
Tamra: 5/6-carboxy-tetramethyl-rhodamine succinimidyl ester

## Acknowledgements

We thank the IBV Imaging Center, University of Oslo for help and expertise in confocal microscopy, and Professor Lucy Robertson for help during the writing process.

## Author contributions

Conceptualization, H.K.G. and T.M.H.; Investigation, H.K.G., R.F., K.H., I.T., E.AW, T.M.H.; Writing, H.K.G., T.M.H., F.A.W., R.F., and K.H.; Project administration, H.K.G.; Funding acquisition, F.A.W. and T.M.H.

## Supplementary information

### Video S1, related to Figure 3

Imaging of Ca^2+^ uncaging experiment showing propagation of Ca^2+^ signal through extension.

### Video S2, related to Figure 7B

Time-lapse video of primary culture during the first 17 hours after seeding. Frames are taken every 3 minutes. For scale bar and other details, see Figure 6.

### Video S3, related to Figure 9

Imaging of Ca^2+^ uncaging experiment showing propagation of Ca^2+^ signal across soma-soma border.

